# Molecular context of pathogenic variants is associated with phenotype and treatment response in SCN8A-related disorders

**DOI:** 10.64898/2026.09.22.753542

**Authors:** Joshua B. Hack, Joseph C Watkins, Michael F Hammer

## Abstract

**Objectives:** Genotype–phenotype studies in rare epilepsies typically relate a pathogenic DNA sequence change to clinical outcome, without considering the broader molecular context of a given variant. Here we ask whether clinical heterogeneity in SCN8A-related disorders (SCN8A-RD) is patterned along molecular dimensions that go beyond the specific genetic alteration, to include position in the linear channel topology, proximity to post-translational modification (PTM) and structural sites, and predicted effects on mRNA splicing. Using the International SCN8A Registry, we map these features against seizure, developmental, and treatment-response phenotypes to identify the regions and molecular feature classes with which clinical variability is associated.

**Methods:** We identify “hot-spots” across the coding sequence for seizure types and severity of developmental disability (DD). We then test for enrichment of clinical features near sites related to PTM and protein structure, identify variants that may alter mRNA splicing, and evaluate the interaction between these features and clinical subgroups.

**Results:** We expand on previous work identifying regions of coding DNA sequence that are pathogenic or benign “hot-spots”. We show regional enrichment of numerous seizure types and severities of DD. Enrichment analysis of PTM and protein structural sites shows distinct enrichment profiles for developmental, seizure, and medication-response features. Finally, we identify 14 variants that are likely to alter RNA splicing.

**Significance:** We provide a higher-resolution map of pathogenic variation across SCN8A and show that seizure types, developmental severity, and medication response are regionally organized along the coding sequence. Proximity to structural and glycosylation sites distinguishes phenotypic subgroups is associated with differential drug response, including a glycosylation–gabapentin relationship, and 14 missense variants are flagged as likely splice-altering, nominating targets for splice-directed therapy and providing candidates for experimental validation of dual pathogenic mechanisms.

**Key Points:**

- Pathogenic variants associated with SCN8A-Related Disorders are regionally enriched and cluster in pathogenic or benign “hot spots” that may inform the classification of variants of uncertain significance.
- Molecular and structural landmarks in *SCN8A* associate with different phenotypic and therapeutic profiles, providing potential insight into developmental prognosis, treatment selection, and underlying seizure mechanism.
- Some pathogenic missense variants may result in aberrant RNA splicing, causing dual mechanisms of pathogenicity. Heterogeneity between patients with identical variants may be explained by differential splicing behavior.

## Introduction

Genotype-phenotype associations are among the most prioritized areas of research in rare diseases, including in childhood epilepsies caused by pathogenic variants in brain expressed voltage-gated sodium channels (Na_V_s). These channels have a common architecture with 4 major domains, each with 6 segments connected by a series of linkers and loops^1^. The four major brain expressed Na_v_s include Na_v_1.1, Na_v_1.2, Na_v_1.3, and Na_v_1.6 encoded by SCN1A, SCN2A, SCN3A, and SCN8A, respectively^2^. The ability to predict clinical outcomes based on the position of a particular variant within this topology is the focus of many studies^3,4^, which have been limited by small cohort sizes. A number of genotype-phenotype studies have been carried out on cohorts with SCN8A-related disorders (SCN8A-RD)^5–8^, which have relied on comparisons of patients carrying the same highly recurrent variant to establish meaningful associations with clinical outcomes^9^. These phenotype-phenotype associations have supported models to predict outcomes and to classify patients into clinically distinct phenotypic subgroups^10,11^. To date, variants under study have been limited to those causing missense, premature termination, frameshift, and splice site alterations.

Consideration of how variants may affect other molecular features of the gene is missing from many of these studies. Post-translational modifications (PTM) contribute substantially to protein function^12,13^. For example PTM of other voltage-gated sodium channels has been shown to alter protein expression in chronic pain syndromes^14^. Phosphorylation, ubiquitylation, and methylation are the most prominent PTM in SCN1A, SCN2A, SCN3A, and SCN8A, each of which can alter expression and function^15^. A further molecular feature that remains largely unexplored is the effect of missense variants on splicing elements ^15^. While variants near canonical splice sites and intronic variants that impact splicing are often addressed, missense and synonymous variants that are distant from canonical splice sites are known to impact splicing products^16^. Overlooking these molecular features restricts our understanding of the possible impacts of each variant and hinders effective clinical management.

To begin to address these gaps, we use the International SCN8A Registry ^17^, PTM and structural annotation databases, and machine-learning predictors of splice-altering behavior to extend genotype–phenotype analysis beyond the amino-acid substitution. We first update the topological map of pathogenic and benign variation across SCN8A at higher resolution than previously achievable ^18^ and show that clinical features, seizure types, developmental severity, and medication response, are themselves regionally distributed across this map, identifying regions in which specific phenotypes are over-represented. We then ask what is molecularly distinctive about these regions, testing whether proximity to PTM, structural, and binding sites, and predicted splice-altering behavior, are associated with distinct clinical and treatment-response profiles. Finally, we relate these molecular features to established clinical subgroups^9^. These analyses cover only a subset of potential consequences; the biophysical and three-dimensional structural mechanisms that contribute to channel dysfunction are not addressed. Within these bounds, our aim is to identify molecular dimensions that are largely unconsidered in prior SCN8A-RD studies and to generate mechanistic and potentially actionable hypotheses, from medication selection to splice-directed therapy, for a heterogeneous population.

## Methods

### Dataset Construction

SCN8A missense variants were curated as pathogenic or benign. Pathogenic variants were identified through literature review, caregiver community engagement, and the International SCN8A Registry^17^ (879 individuals, 420 unique missense variants; **Figure S1**). Benign variants were drawn from gnomAD v4.1.1^19^ retaining only those classified “benign” or “likely benign” by ACMG criteria^20^. Pathogenic and benign variants were deduplicated by coding sequence (CDS) position, yielding 381 and 767 unique coding DNA sequence (CDS) positions, respectively.

Caregiver-reported data from the International SCN8A Registry were collected for 511 patients with genetically confirmed SCN8A variants. Seizure semiology and 25 developmental milestones across four domains (gross motor, fine motor, social motor, language) were reported by caregivers, with social motor further split into adaptive/self-care and social/communication subdomains. Developmental quotient (DQ) was calculated as the neurotypical age of skill acquisition divided by the age of the patient at assessment, multiplied by 100. Patients were assigned to a clinical subgroup defined by Hack et al.^9^ using machine learning with manual validation. The study population is described in **Table S1**.

Each variant was annotated with seven splice-effect predictors (SpliceAI^21^, Pangolin^22^, MMSplice^23^, MaxEntScan^24^, SQUIRLS^25^, SPiP^26^, and dbscSNV^27^), and PTM sites were compiled from UniProt^28^, dbPTM^29^, and PhosphoSitePlus^30^; variants were flagged as within 50 nucleotides (nt) or 100 nt of a PTM site. The final dataset includes 386 patient profiles with at least partial data for seizure onset and semiology, developmental milestones, distance from PTM, predicted splice-altering status, and clinical subgroup.

### Genetic Topology

Cumulative distribution curves of pathogenic and benign variants across the SCN8A coding sequence (5,940 nt) were used to define pathogenic and benign “hot spots” as described previously ^18,31^ (see Supplementary Methods for the region-classification procedure). Anderson–Darling tests compared the pathogenic, benign, and uniform distributions; chi-square and two-sided Fisher’s exact tests assessed per-segment enrichment (significant at Storey’s q<0.05^32^).

### Symptom Topology

DQ was stratified into profound DD (DQ≤30), severe DD (30<DQ≤50), moderate DD (50<DQ≤70), and mild DD/neurotypical (DQ>70). Cumulative distribution curves evaluated DD-severity categories and seizure types across the coding sequence. For sliding-window enrichment, a 300-nt window was advanced in 50-nt steps; at each window a 2×2 Fisher’s exact test compared feature-positive versus feature-negative patients inside versus outside the window, with p-values expressed as signed −log₁₀(p). Storey’s q-values were computed across all sliding-window tests combined.

### Post-Translational Modification Site Enrichment

PTM sites were categorized as phosphorylation, glycosylation, binding, structurally relevant sites, and disordered regions. Windows of 50 nt and 100 nt on either side of each site were tested for symptom enrichment by two-sided Fisher’s exact tests. Enrichment was reported as −log₁₀(p) (radar plots, by seizure type and developmental domain) and |log₂(OR)| (lollipop plots, 50-nt windows); throughout, p<0.05 was treated as nominally significant and exploratory (hypothesis-generating).

### Splice-Altering Variants

Per-tool thresholds were set from the literature to achieve 95% specificity, classifying each variant as likely splice-altering or normal (see Supplementary Methods). A variant was classified splice-altering by the composite if both SpliceAI and Pangolin flagged it and at least one additional line of evidence was present: disruption of a canonical splice site (MaxEntScan 5′ or 3′) or flagging by ≥2 of five ensemble/conservation tools (MMSplice, SPiP, dbscSNV, SQUIRLS). Symptom enrichment was tested per tool by two-sided Fisher’s exact tests with 1,000-replicate individual-level bootstrapping, yielding bootstrapped odds ratios and confidence intervals (**Table S4**). A final list of predicted splice-altering variants was generated for future validation.

### Similarity Analysis of Splice-altering versus Recurrent Variants

Likely splice-altering variants were compared to recurrent variants (>4 occurrences) and to variants within 50 nt of a PTM. For each variant, a clinical profile vector was built from the proportion of its carriers exhibiting each feature (seizure types, DQ, and developmental milestones). Pairwise cosine similarity was computed between profile vectors, and variants were clustered (Ward’s method), ordinated by PCA, and embedded by UMAP (parameters in Supplementary Methods).

### Clinical Subgroup Analysis

Among the 226 patients with an assigned subgroup, each of the five subgroups was compared with all others pooled using two-sided Fisher’s exact tests (significant at q<0.05). Symptom-category radar plots were constructed as above, with an additional radar plot using PTM categories and splice-altering status as spokes; per-subgroup lollipop plots showed enrichment of splice-altering variants and variants within 50 nt of a PTM (nominal p<0.05, hypothesis-generating).

## Results

### Pathogenic SCN8A Variants are Topologically Enriched

Of the 49 unique segments in Na_V_1.6, 13 showed statistically significant enrichment or depletion of pathogenic SCN8A variants (**Figure 1A**). Eleven of these segments were enriched and the DI-DII and DII-DIII cytoplasmic loops were depleted (**Table S2**); the DIII–DIV cytoplasmic loop, by contrast, was enriched. Regions depleted in pathogenic variants are generally inter-segment linkers or early segments of each domain, while regions enriched for pathogenic variants are generally segments 4–6 in each domain (**Table S2**).

**Figure 1.**
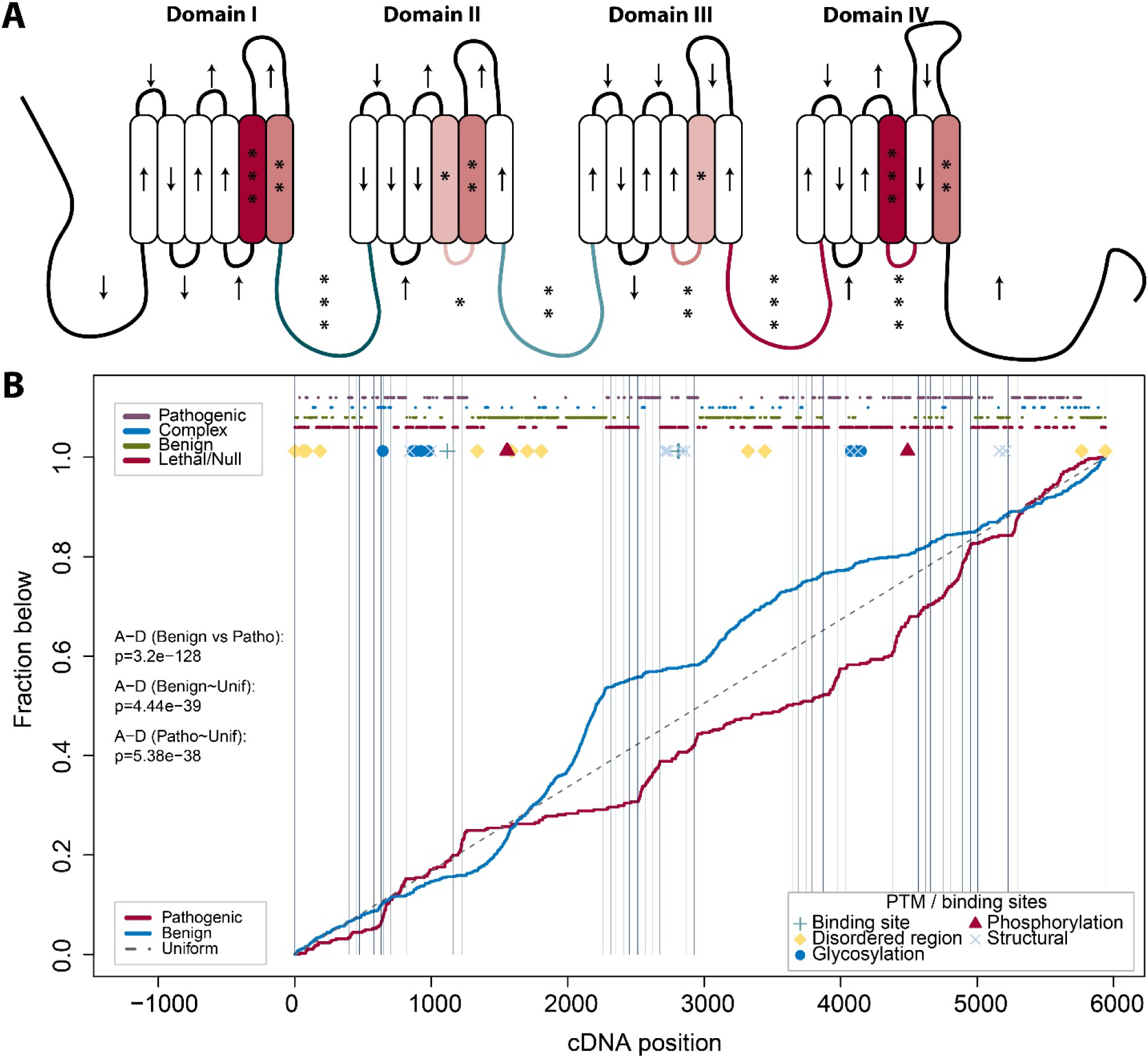
Topology of pathogenic and benign SCN8A variants. A) Enrichment (*red*) or depletion (*blue*) of pathogenic variants by segment of Na_V_1.6. Storey’s q-value is marked by asterisks for q<0.05 (*), q<0.01 (**), and q<0.001 (***). Segments without significant finding are marked with arrows showing enrichment (up) or depletion (down). B) Cumulative distribution of variants reported to be pathogenic (*red*) or benign (*blue*) across the coding sequence of SCN8A. Anderson-Darling test results are shown comparing the distributions of benign versus pathogenic, benign versus uniform distribution, and pathogenic versus uniform distribution. Vertical lines indicate boundaries between segments.

Neither pathogenic (Anderson-Darling p-value = 5.4x10^-38^) nor benign (p-value = 4.4x10^-39^) variants are uniformly distributed across the coding sequence (**Figure 1B**). The first two cytoplasmic loops exhibit long stretches of benign variants, while the DIII–DIV loop is predominantly pathogenic (**Figure 1B**). A greater portion of the C-terminus is enriched for pathogenic variants, while the N-terminus splits into benign or lethal regions. Disordered regions show the highest tolerance to variation, while binding sites, disulfide bridges, and glycosylation sites are embedded in long stretches of lethal or pathogenic regions (**Figure 1B**). The three phosphorylation sites appear to have differential tolerance to variation.

### Clinical features are topologically enriched

Analysis of the distribution of clinical features such as seizure types at initial onset and extent of DD reveals non-uniform distribution of the seizures across the coding sequence (**Supplementary Data 1**). Myoclonic seizures appear to be enriched in Domains I and II, while focal seizures appear to be more prominent in Domains III and IV (**Figure 2A**). Sliding-window analysis identifies regional enrichment of infantile spasms (IS), myoclonic, tonic, epileptic spasms (ES) (i.e., spasms occurring beyond 2 years old), absence, bilateral tonic-clonic (BTC), atonic, and clonic (**Figure 2B; Supplementary Data 1**). Only IS, tonic, and myoclonic seizures show stretches of reduced occurrence that reached statistical significance.

**Figure 2.**
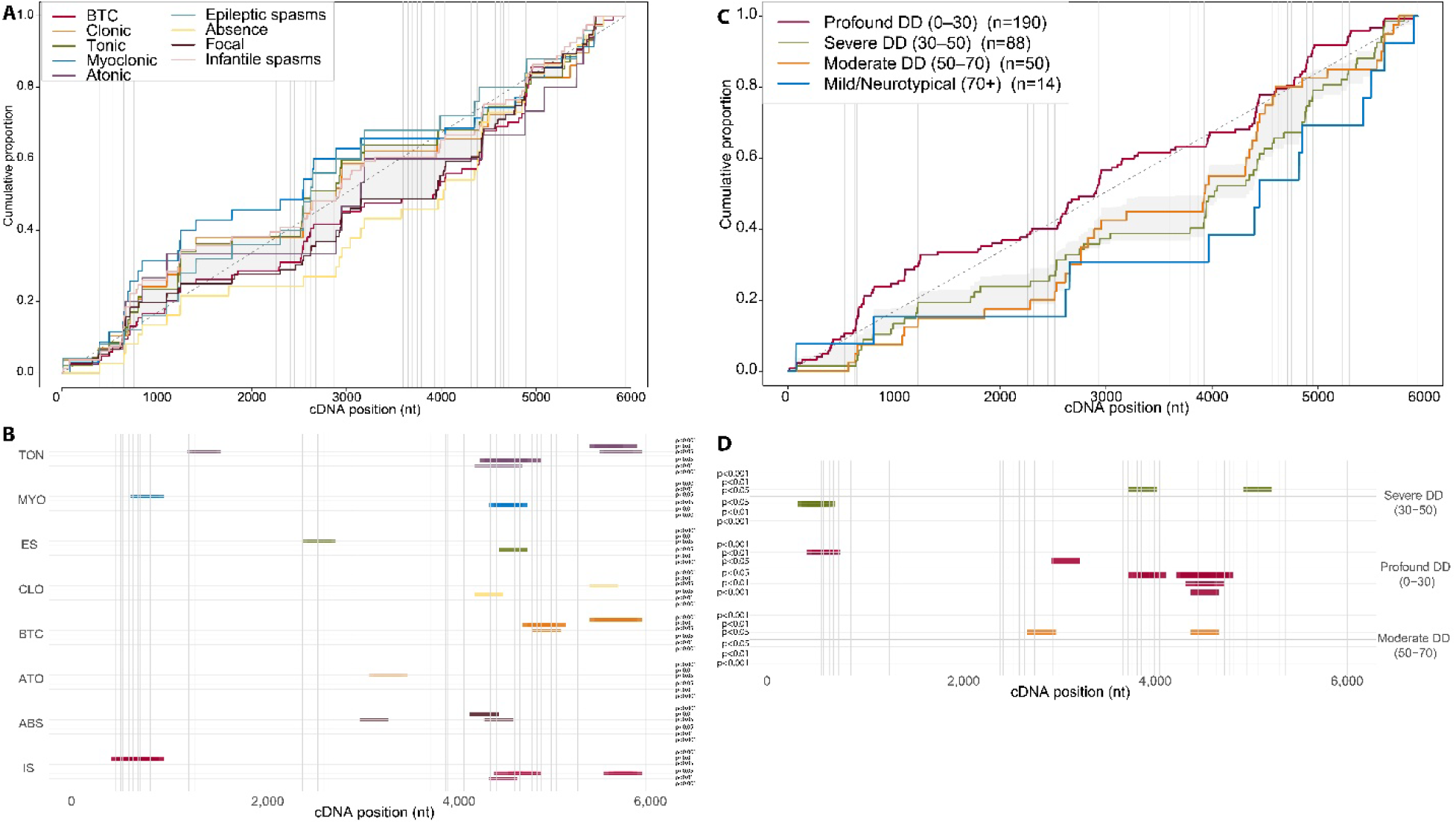
Distribution of clinical features across the coding sequence. A) Cumulative distribution curves of patients grouped by seizure type against a uniform reference (*dashed*). B) Sliding window enrichment analysis using 300 nucleotide long windows with 50 nucleotide steps for seizure types. Windows above midline for each feature indicate enrichment; windows below midline indicate depletion. Vertical lines mark regional boundaries. C) Cumulative distribution curves of patients grouped by developmental delay (DD) severity based on developmental quotient (DQ): profound DD (0-30), severe DD (30-50), moderate DD (50-70), and mild DD and neurotypical (70+). D) Sliding window enrichment for DD severity groups. BTC: Bilateral Tonic-Clonic; CLO: Clonic; TON: Tonic; MYO: Myoclonic; ATO: Atonic; ES: Epileptic Spasms; ABS: Absence; FOC: Focal; IS: Infantile Spasms; DD: Developmental Delay.

Severity of developmental delay is also distributed differently from the underlying variant distribution (**Figure 2C**). The majority of the population presented with profound DD and few presented with mild DD or neurotypical development (n=14). Severity of DD was regionally structured (**Figure 2D**): profound DD was enriched in early Domain I and depleted around p.Arg1475, severe DD was depleted in early Domain I, and moderate DD was enriched around p.Arg1475. A finer-scale analysis (150 nt windows, 25 nt steps) additionally revealed enrichment of mild DD/neurotypical development from 5351–5575 nt (**Figure S2**).

### Symptom Association with Modification and Structural Sites

Clinical profiles across sites relevant for protein structure (disordered regions, disulfide bridges), function (binding sites), and post-translational modification (phosphorylation and glycosylation) are broadly similar, particularly for developmental milestones (**Figure 3A-F**). Patients with variants within 50 nt of structural sites had reduced success on non–sodium-channel-blocker regimens and fewer BTC seizures, and were enriched for standing without support, naming colors, and stacking blocks (**Figure 3G**). Patients with variants within 50nt of a binding site were depleted for developing laughing or vocalizing (**Figure 3H**). Phosphorylation sites showed enrichment for later-acquired developmental milestones: reading, brushing teeth, and speaking in full phrases (**Figure 3I**). Glycosylation sites were enriched for IS, gabapentin success, and diazepam success while being depleted in babbling, lacosamide success, carbamazepine success, and sitting (**Figure 3J**). All odds ratios and p-values are presented in **Table S3**. Intrinsically disordered regions yielded no significant results at 50nt windows but were enriched for IS when the window was expanded to 100nt (**Table S3**).

**Figure 3.**
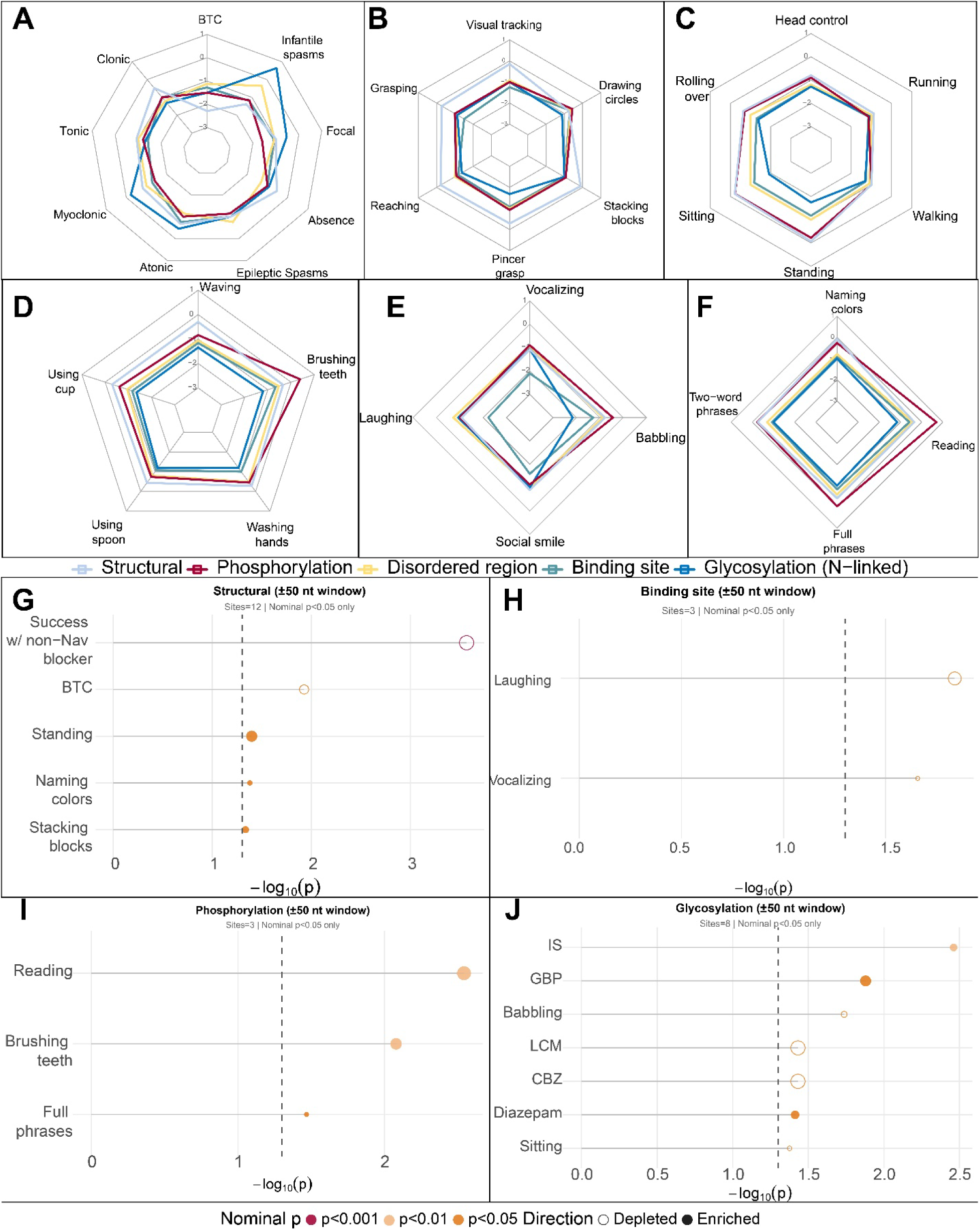
Enrichment of symptoms based on proximity to post-translational modification sites and structurally relevant features. Radar plots showing enrichment of symptoms within 50 nucleotides of a post-translational modification site or structurally relevant feature. Spokes represent distinct clinical features; values are signed -log_10_(p) from two-sided Fisher’s exact tests comparing the proximity to PTM sites of feature-positive and feature-negative patients. Panels show (A) seizure types, (B) fine motor milestones, (C) gross motor milestones, (D) adaptive/self-care milestones, (E) social/communication milestones, and (F) language milestones. (G-J) Lollipop plots of nominally significant (p<0.05) enrichment within 50 nucleotides of structural sites, binding sites, phosphorylation, and glycosylation sites. Points are colored by significance tier (p < 0.001, dark red; p < 0.01, orange-red; p < 0.05, orange) and shaped by direction (filled = enriched, open = depleted). Point size reflects |log₂(OR)|.

### Missense Variants in SCN8A are Predicted to Alter Splicing

SpliceAI and Pangolin, the most similar in construction, diverged in their associations: both were depleted for reading, but only Pangolin was enriched for carbamazepine success (**Figure 4A**). The full set of per-tool enrichments and depletions is given in Figure 4A and Table S4. By the composite flag, predicted splice-altering variants were enriched for carbamazepine success and depleted for speaking in phrases and reading (**Figure 4B**). dbscSNV was excluded because few variants had predictions available..

**Figure 4.**
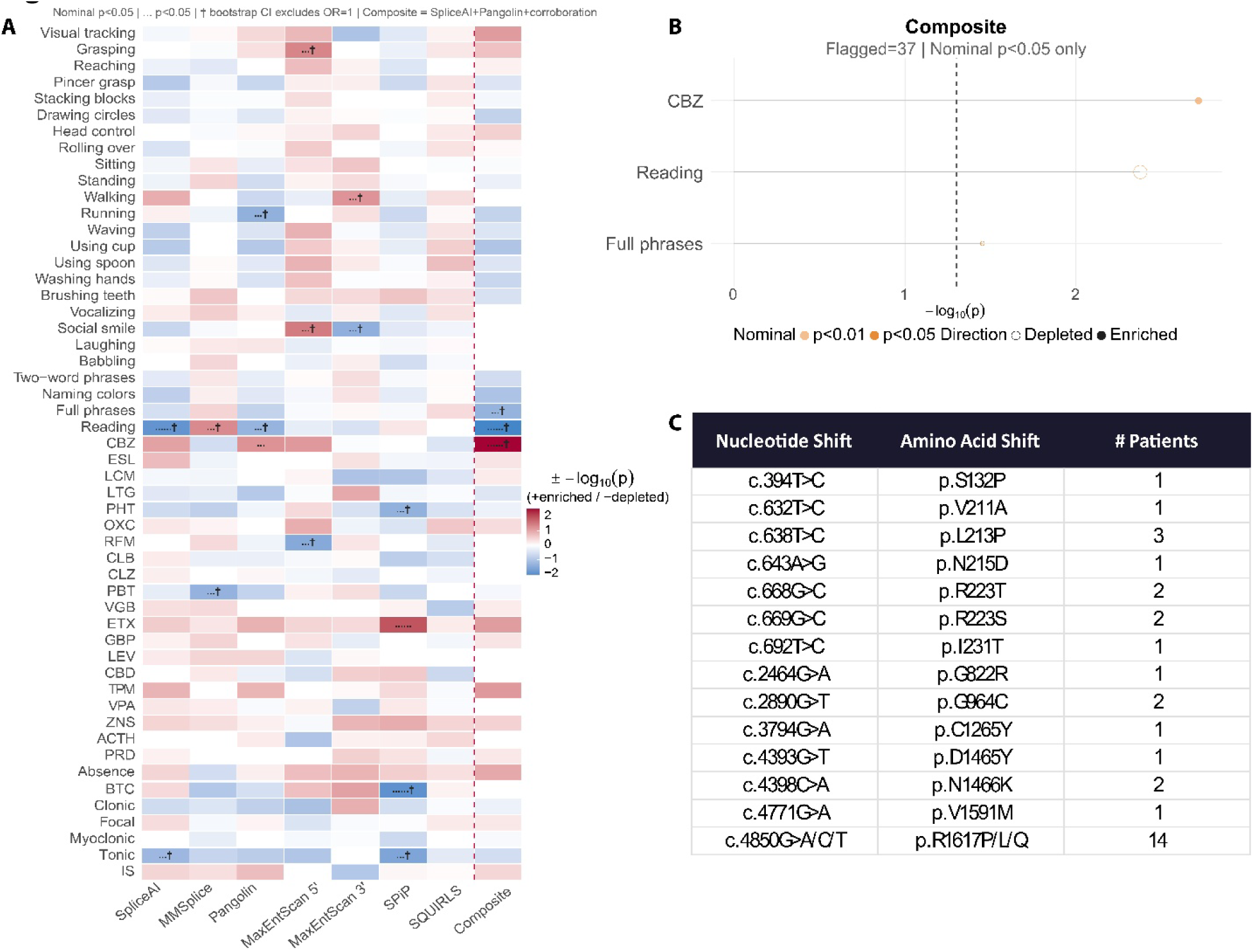
Symptom enrichment in variants predicted to be splice-altering by composite. A) Heatmap of symptom enrichment for variants flagged by each of 6 tools and the composite decision. Color encodes signed -log₁₀(p) from two-sided Fisher’s exact tests (flagged vs. below-threshold patients); red indicates enrichment in flagged patients, blue indicates depletion. † bootstrap 95% CI for the OR excludes 1. dbscSNV was excluded because few variants had dbscSNV predictions available. B) Lollipop plot of nominally significant enrichments for the composite classification. CBZ= carbamazepine beneficial. C) Table of composite-flagged variants in this cohort. Number of patients includes all patients reported in literature, in the Registry, and collected through community interaction.

Fourteen variants met the criteria to be flagged as likely splice-altering (**Figure 4C**). Six recur, with two affecting the same codon (c.668 and c.669, both affecting p.223), and the set includes the recurrent p.R1617Q/L/P, which has a heterogeneous clinical presentation.

### Clinical profiles of splice-altering variants resemble those near glycosylation and structural sites

Variant-level Pearson correlation analysis shows that likely splice-altering variants correlate negatively with BTC seizures and positively with atonic seizures, while variants near phosphorylation sites correlate positively with gross motor development (**Figure 5A, Table S5**). Hierarchical clustering shows splice-altering variants are most similar to variants close to glycosylation and structural sites (**Figure 5B**). The first two principal components account for 64.5% of the variance, and both the principal component analysis and the UMAP embedding reproduce the clustering result that the two recurrent splice-altering variants, p.223 and p.1617, present distinctly from each other (**Figure 5C-D**).

**Figure 5.**
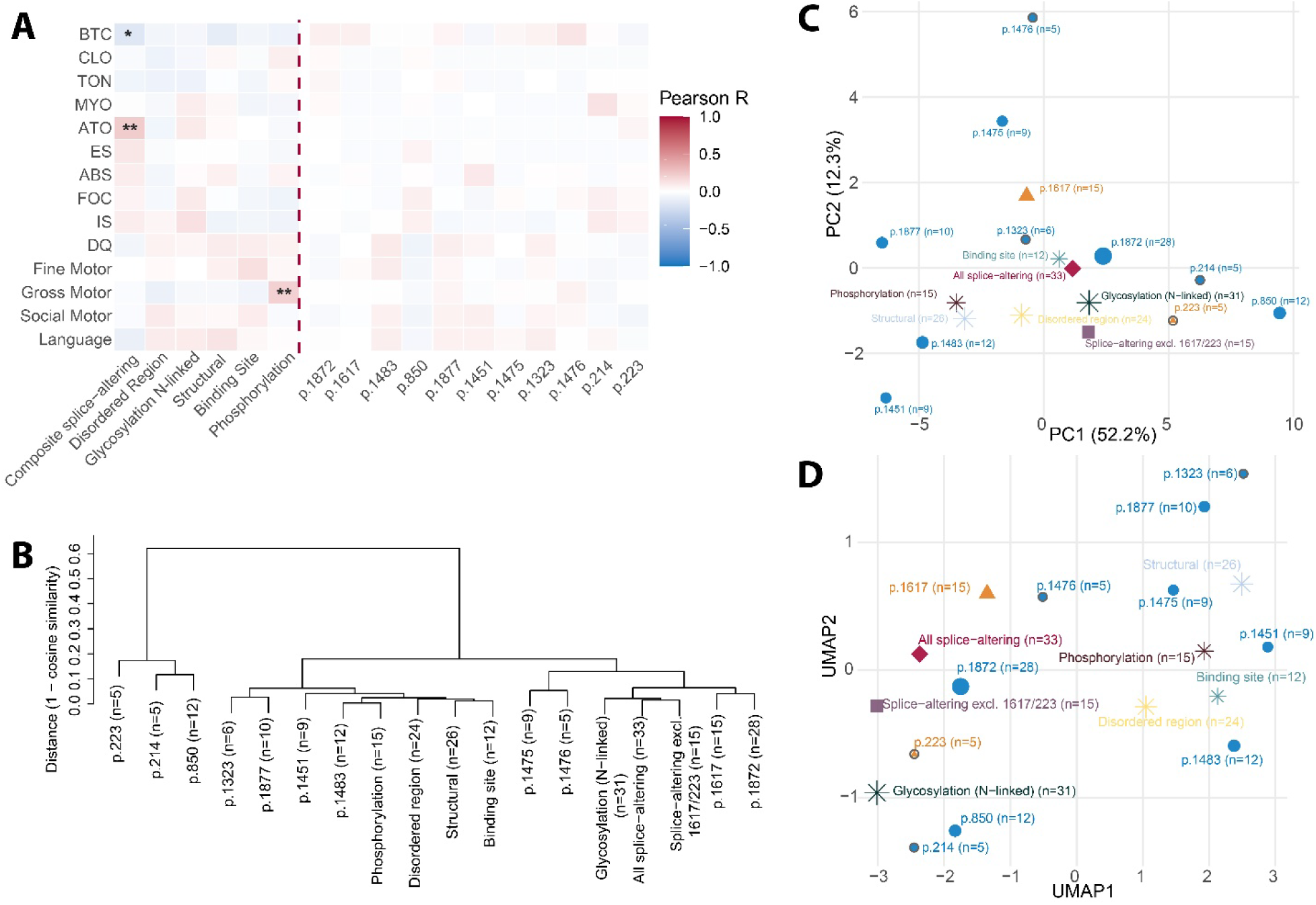
Similarity of clinical profiles among highly recurrent variants and predicted splice-altering variants. A) Variant-level Pearson correlation matrix between seizure type and developmental quotient and molecularly relevant features and recurrent variants. *: p<0.05; **: p<0.01. BTC: Bilateral Tonic-Clonic; CLO: Clonic; TON: Tonic; MYO: Myoclonic; ATO: Atonic; ES: Epileptic Spasms; ABS: Absence; FOC: Focal; IS: Infantile Spasms; DQ: Developmental Quotient. B) Hierarchical clustering dendrogram of recurrent SCN8A variants (n>4) and the splice-altering class of variants using Ward’s method applied to cosine distance of clinical profile vectors, determined by seizure types and developmental milestones acquired. Two groups of splice-altering variants are presented: one including the recurrent p.223 and p.1617 variants and one excluding these. C) Principal component analysis of clinical profile vectors. Points are sized by number of patients with each variant. D) UMAP of clinical profile vectors. Open circles denote variants with fewer than 8 patients.

### Clinical subgroups associate with different features

Across the five subgroups defined by Hack et al.^9^ (LOF with seizures (LOF+), LOF without seizures (LOF−), GOF-DE, GOF-EE, and GOF-DEE) enrichment analysis showed enrichment of IS (OR=3.07, p=7.1×10⁻⁴) in the DEE subgroup (**Figure 6A**). When considering protein and molecular landmarks, patients in the LOF-subgroup were enriched within 50nt of structural components (OR=4.1, p=8.0x10^-3^) and 100nt of binding sites (OR=3.6, p=0.045), while the DE subgroup was depleted within 100nt of glycosylation sites (OR=0.13, p=0.02) (**Figure 6B**). The LOF− subgroup was also enriched for many developmental skills across domains (**Figure 6C–G**).

**Figure 6.**
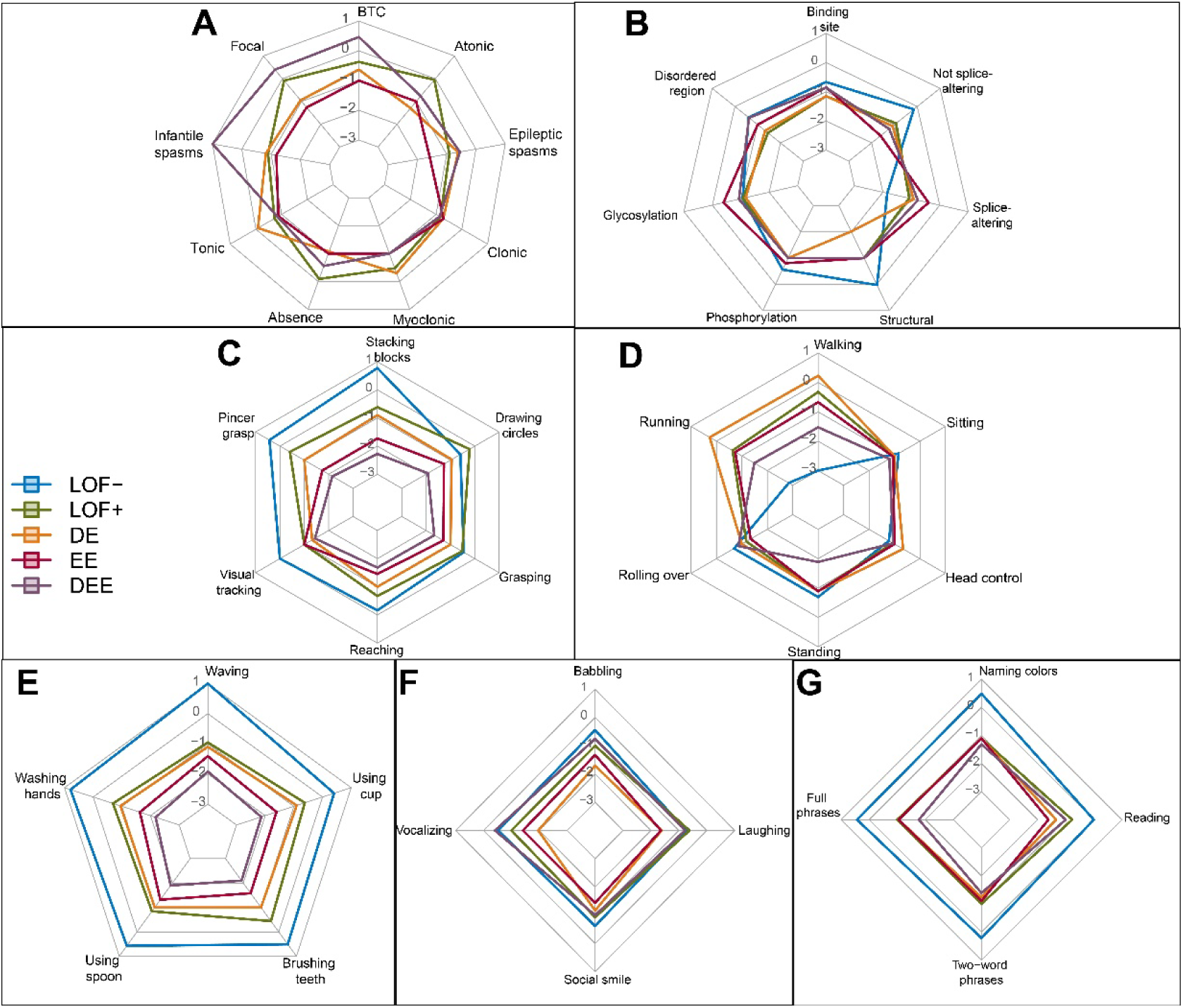
Symptom profiles across clinical subgroups. Radar plots showing enrichment of clinical features within each of five subgroups: LOF-, LOF+, GOF-DE, GOF-EE, and GOF-DEE. Values are signed -log_10_(p) values from two-sided Fisher’s exact tests. A) Seizure types (excluding LOF-). B) Molecular features including predicted splice-altering, predicted non-splice-altering, phosphorylation sites, glycosylation sites, and structurally relevant features. C) Fine motor developmental milestones. D) Gross motor developmental milestones. E) Adaptive/self-care milestones. F) Social/communication milestones. G) Language milestones.

## Discussion

Identifying meaningful genotype-phenotype correlations in rare disease is constrained by small populations, clinical heterogeneity within the same variant, and the diversity of causative variants; outside of highly recurrent variants, predicting an individual’s natural history remains difficult. This study addresses these barriers by placing pathogenic variants within the context of the molecular architecture of Na_V_1.6 and asking how clinical features are distributed across that architecture. We first establish an updated topological map of a large set of known pathogenic variants (scn8a.net), then show that phenotypes are themselves regionally organized upon it. Finally, we ask whether variants in proximity to PTM and structural sites and in predicted splicing consequences are associated with specific clinical features or phenotypic subgroups.

### An updated pathogenicity map

The method used to identify “hot-spots” of pathogenicity was introduced in Encinas et al.^18^ Since 2020, the catalog of both benign and pathogenic SCN8A variants has expanded substantially, permitting higher resolution in locating where these hot spots reside. Where Encinas et al.^18^ identified only eight pathogenic regions amid large benign stretches, the present map expands the pathogenic regions while shrinking those deemed to be benign. The increased resolution also raises the number of “lethal” regions in which no variant is reported. Importantly, these regions are not necessarily lethal but may harbor unreported, tolerated variants (or be associated with yet to be described neurological phenotypes). Further analysis of evolutionary conservation and predicted amino acid effects at these sites will clarify which regions are truly intolerant.

This topology also aids variant classification in that it provides evidence for classifying novel variants as likely pathogenic or benign^31^, and the updated map offers an additional line of evidence for SCN8A variants of uncertain significance. A limitation of the original topology was the scarcity of variants associated with SCN8A presenting without epilepsy. Since 2020, reports of SCN8A variants presenting primarily with autistic features and little or no epilepsy have increased, allowing for the current map to capture a broader phenotypic landscape^18^. The higher density map further refines the complex and lethal regions as more such variants are reported.

At the segment level, our results reproduce the enrichment and depletion trends reported by Encinas et al.^18^, with only minor discrepancies in magnitude and none in direction. The disparate tolerance of the N- and C-termini and of the DIII-DIV loop relative to the other intracellular loops remains notable. Pathogenicity of DIII-DIV loop variants is expected since this loop functions as the inactivation gate of Na_V_1.6. Enrichment of the C-terminus, hypothesized by Encinas et al. to reflect its interaction with calmodulin, may additionally involve protein-protein interactions with fibroblast growth factor homologous factors (FHFs) which bind to the proximal globular EF-hand-like domain and disruption of palmitoylation at Cys1978^33–35^. The higher resolution here localizes most C-terminal hot spots to the proximal C-terminus and suggests variation at Cys1978 may be tolerated; confirmation of the calmodulin and FHF mechanisms awaits further work.

### Phenotypes are regionally organized on the map

Beyond pathogenicity, the sliding-window scan identifies stretches of the coding sequence in which specific clinical features are over-represented. Infantile spasms (IS) and epileptic spasms (ES), among the more detrimental seizure types in SCN8A-RD given their impact on developmental outcomes^8^, are enriched early (p.133-p.300) in Domain I and around p.Arg850Gln (p.783-p.900). While the variants in DI are enriched for IS, p.Arg850Gln is enriched for ES (i.e., spasms that persist beyond infancy). Myoclonic seizures show a similar topology to IS, consistent with a shared regional distribution of these seizure types. Absence seizures are not only enriched around highly recurrent loss of function (LOF) variants, but also early in the DII-DIII Loop. Absence seizures are among the primary features of individuals carrying variants with LOF effects^5,10^, consistent with a concentration of LOF-associated positions early in the intracellular loops.

Developmental severity is similarly regionalized. A substantial share of the SCN8A population experiences profound DD, a phenotype that displays strong enrichment and depletion in particular regions. The risk of profound DD appears to be highest in the early-DI and lowest around p.Arg1475, the latter being most strongly enriched for moderate DD. Despite the enrichment for profound DD, the early-DI is depleted for Severe DD, suggesting that patients with variants in this stretch rarely achieve more than a few developmental milestones.

### Structural sites, loss of function, and medication response

To our knowledge, the association of post-translational modification and structural sites with phenotype has been largely unconsidered in prior SCN8A-RD studies. Because PTM sites are highly conserved, variants near them are more likely to be pathogenic^36^. Most sites within the “structural” category are disulfide bridges, which are most strongly associated with the LOF-without-seizures (LOF−) subgroup. Because these bridges are critical for protein folding, nearby variants may interrupt protein trafficking or lead to failure of the channel to reach the membrane, thus reducing ion current (i.e., loss of function). Proximity to these structural sites is also the only association within 50nt of a modification or structural site to survive multiple-testing correction, appearing as a depletion for success on non–sodium-channel-blocker regimens. This links medication response to the structural/LOF axis identifies a candidate criterion for treatment selection, though the direction of the relationship warrants careful interpretation and is a priority for follow-up.

Phosphorylation sites in Na_V_ channels are highly conserved across channel subtypes and differentially regulate channel function in Na_V_1.1, Na_V_1.2, and Na_V_1.6, with phosphorylation of NaV1.6 either enhancing or reducing channel activity depending on the kinase involved^37^. In Na_V_1.1 and Na_V_1.2, phosphorylation can reduce channel function, providing a possible target to compensate for GOF variants^37,38^. Variants at these sites may therefore disrupt channel regulation through a mechanism not addressed by conventional therapies, and therapies that modulate phosphorylation may compensate for either GOF or LOF effects across more than one Na_V_ channel.

### Glycosylation and gabapentin response

Glycosylation sites are of particular interest due to their enrichment for IS and for success on GPB, while being depleted for success on LCM and CBZ (**Figure 3J**). Gabapentin binds the α2δ auxiliary subunit and interacts with its highly N-glycosylated regions ^39–41^. This offers a plausible mechanism for its efficacy in this subset despite limited efficacy across the SCN8A-RD population overall ^42^, and constitutes a directly testable pharmacological hypothesis^41^.

### Splice-altering missense variants and therapeutic implications

A frequently overlooked class comprises splice-altering missense variants that lie outside canonical splice sites. Despite a long history of splice-prediction tools, these have rarely been applied to channelopathies to investigate mechanism. This study provides a short list of variants that should be considered likely splice-altering and that warrant experimental validation. Both the p.R223T/S and p.R1617P/L/Q are clinically heterogeneous, which may be explained by splice-altering behavior. Notably, both lie deep within exons and may introduce cryptic splice sites, yielding a truncated protein with variability in the proportion of aberrantly spliced product contributing to the range of presentations. These variants also point to alternative therapeutic approaches, for example through anti-sense oligos^43^, exon-specific U1s^44^, or ADAR-mediated editing^45^, that may be required to fully rescue the phenotype rather than relying on anti-seizure medications alone.

### Integration with phenotypic subgroups

Taken together, these molecular dimensions converge on the established phenotypic subgroups^9,46^. Structural- and binding-site proximity is associated with the LOF− subgroup, glycosylation-site proximity is depleted in the GOF-DE subgroup, and the DEE subgroup is enriched for IS. These associations indicate that the LOF/DE/EE/DEE grouping corresponds in part to the distribution of variants relative to these molecular features. Genotype–phenotype associations, such as that between phosphorylation-site proximity and reading, further support interpreting these subgroups as partly reflecting the molecular context of the variant rather than as purely clinical categories.

## Limitations and Further Considerations

While this study generates several hypotheses for the underlying mechanisms of heterogeneity among individuals with SCN8A-RD, several caveats should be considered when interpreting these results. Apart from the pathogenicity map, the segment-level enrichments, and the association within 50 nt of a structural site that survives multiple-testing correction, many associations reported here are nominally significant. This is expected in a rare disease with phenotypic heterogeneity and small cohorts. As such, these findings should be interpreted as preliminary statistical associations rather than fully replicated correlations. Prospective validation will require larger cohorts and/or higher-resolution phenotyping. One practical route to greater power is to aggregate patients across different variants that share a molecular feature (e.g., comparable distance to a PTM site, or shared predicted splice-altering behavior) rather than relying on individual recurrent variants.

The analyses situate each variant along the linear coding sequence and its annotated features, yet do not capture how a variant alters the biophysical properties of NaV1.6^47^, nor how its position in the folded, three-dimensional channel, including proximity to the ion-conducting pore, reshapes function^33,48^. *In vitro* electrophysiological characterization that distinguishes gain-from loss-of-function effects is available for only a minority of reported variants, and structure-guided analysis was beyond the present scope. Classification of variants as gain- or loss-of-function may also oversimplify the electrophysiological impact of a variant, as the independent functional features are spectral and not necessarily concordant. Channel-level consequences are in turn difficult to extrapolate to the neuronal or circuit level. Integrating biophysical and structural dimensions with the molecular topology described here is therefore a priority for future work.

Finally, tools for predicting splice-altering behavior vary considerably, reflecting differences in training data and model design, and agree only partially on which variants are affected^21–27,49,50^. The composite metric used here is a pragmatic compromise that may be overly restrictive, missing true splice-altering variants, or too permissive, flagging false positives. The 14 variants nominated here therefore require experimental validation, and improved tools or consensus metrics are needed to establish which predictors are reliable and under what conditions.

In conclusion, clinical variability in SCN8A-RD is patterned along multiple molecular dimensions of a variant including its topological position and its proximity to modification, structural, and splice-relevant features, beyond the amino-acid substitution alone. Several of these associations carry immediate interpretive and therapeutic implications, from VUS classification to medication selection and candidate splice-directed targets. Considering variants in their complete molecular context should improve both mechanistic understanding and clinical management in SCN8A-RD.

## Data availability statement

All relevant data and code for replication are available at 10.5281/zenodo.21383890. Requests for additional de-identified data will be considered upon request to the corresponding author.

## Ethics approval statement

The study was approved by the University of Arizona Institutional Review Board (#1603487278).

## Patient consent statement

All caregivers of individuals with SCN8A consented to participate prior to completing the questionnaire.

## Permission to reproduce material from other sources

There are no materials from other sources.

