## Supplementary Methods for "Molecular context of pathogenic variants is associated with phenotype and treatment response in SCN8A-related disorders"

Supplementary Materials

### Supplementary Methods

***Genetic topology — region classification***

The distribution curves were split into regions deemed “steep” or “flat” depending on whether there was an excess of variants across a sequence. Regions were classified as null where both distributions were flat, complex if both were steep, pathogenic if the patient distribution was steep while the gnomAD distribution remained flat, or benign if the gnomAD distribution was steep while the patient distribution remained flat. The absence of reported variants in null regions could reflect poor sequence coverage, undiscovered or unreported variants, or intolerance to variation resulting in lethality.

***Splice-altering thresholds***

For each splice-effect predictor, a threshold was set from the literature to meet 95% specificity, and each variant was classified as likely splice-altering or “normal” pathogenic behavior per tool. For analyses yielding p<0.05, additional bootstrapping validation was performed.

***Similarity analysis — parameters***

PCA was performed on the profile matrix with centering and unit-variance scaling. UMAP used a cosine metric, 10 neighbors, and a minimum distance of 0.2. Clustering used Ward’s minimum-variance method on cosine distance.

### Supplementary Figures

***Legends***

**Supplementary Figure 1.** Genetic topology of all pathogenic SCN8A variants used in this study. Yellow stars indicate singletons, orange stars indicate recurrent variants with the number of reported patients carrying a variant at that amino acid position.

**Supplementary Figure 2.** Sliding window enrichment analysis using 150 nucleotide long windows with 25 nucleotide steps for severity of developmental delay. Windows above midline for each feature indicate enrichment; windows below midline indicate depletion. Vertical lines mark regional boundaries.

Supplementary Figure 1.

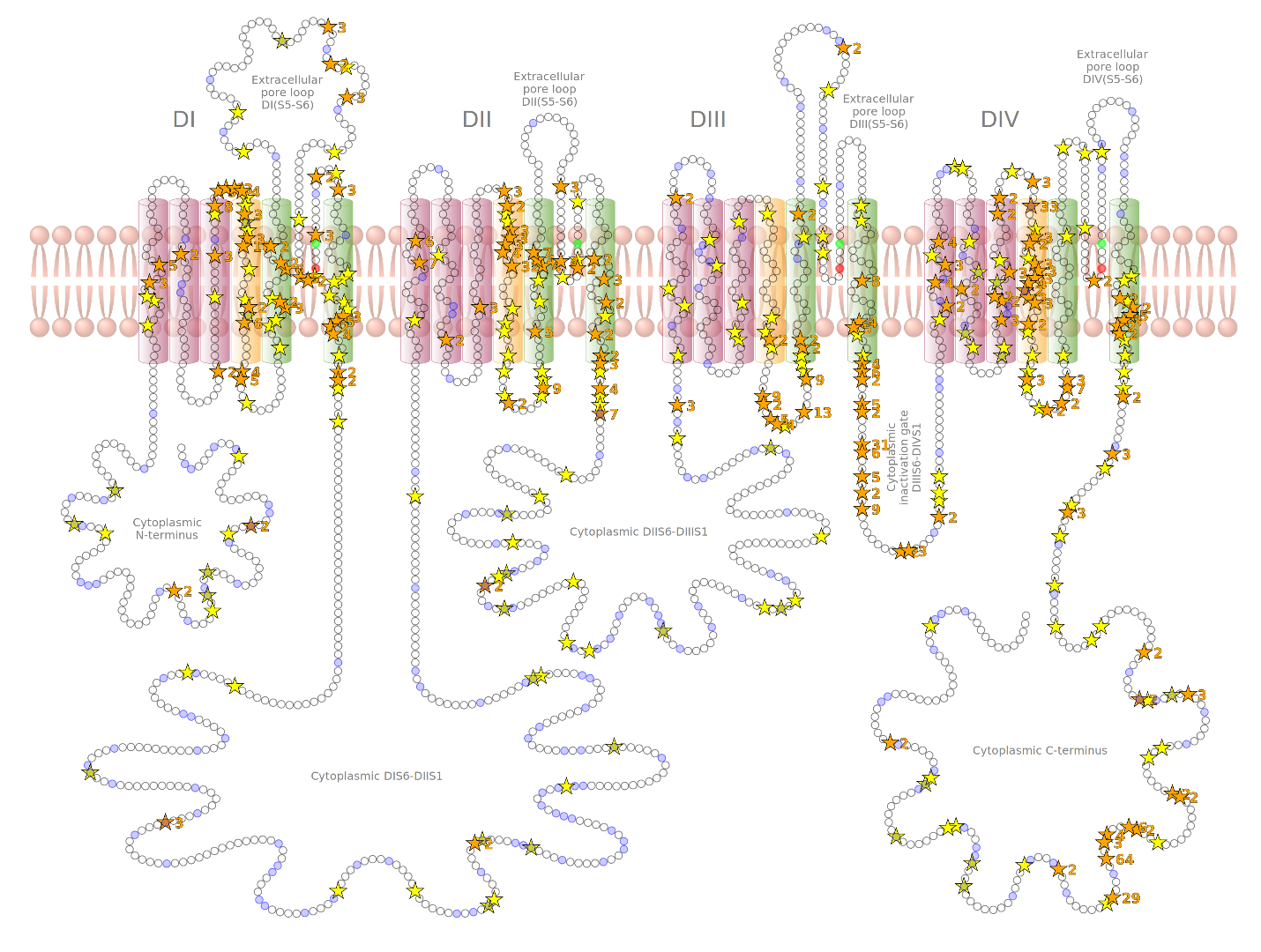

**Supplementary Figure 2.**

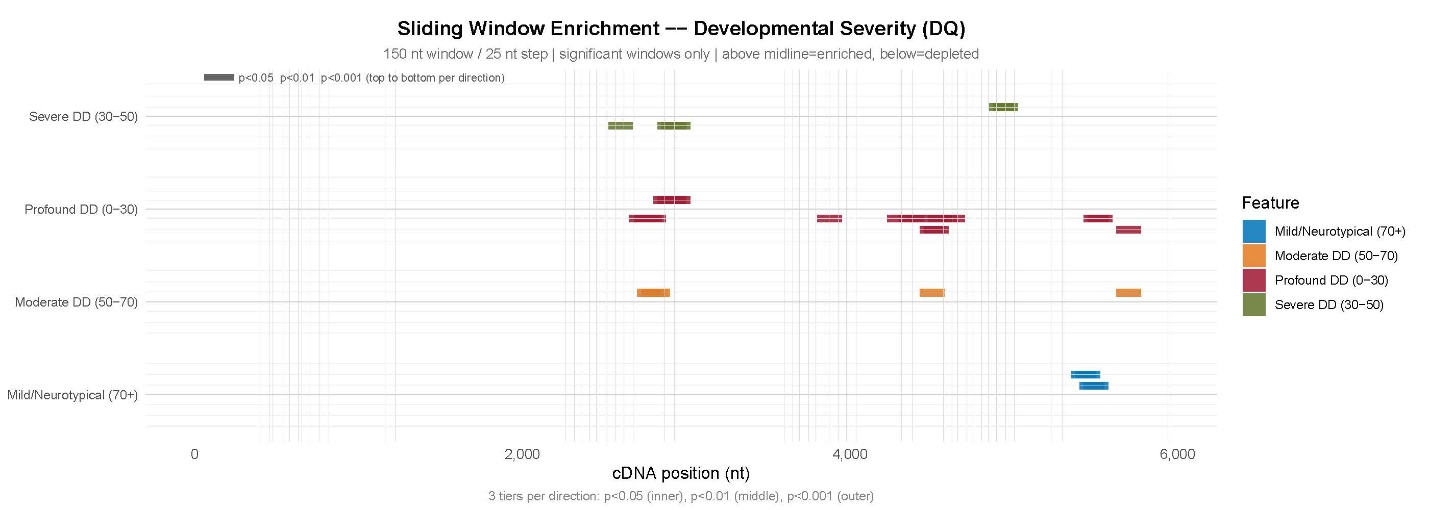

### Supplementary Tables

**Supplementary Table 1.** Study population statistics.

| Characteristic | N (%) or Median (IQR) |
| --- | --- |
| --- DEMOGRAPHICS & ONSET --- |  |
| N (total) | 386 |
| Seizure onset (months) | 4.0 (2.0–8.0) |
| DD onset (months) | 6.0 (2.8–48.0) |
| Neonatal onset | 74 (35.6%) |
| VUS | 116 (30.1%) |
| Loss of Function variant | 44 (24.4%) |
| --- CLINICAL SUBGROUP --- | 226 |
| Loss of Function-No Seizures | 45 (19.9%) |
| Loss of Function-Seizures | 39 (17.3%) |
| Gain of Function- Developmental Encephalopathy | 38 (16.8%) |
| Gain of Function- Epileptic Encephalopathy | 33 (14.6%) |
| Gain of Function- Developmental and Epileptic Encephalopathy | 71 (31.4%) |
| --- SEIZURE TYPES: GENERALISED --- |  |
| Atonic | 17 (4.4%) |
| Bilateral tonic-clonic | 149 (38.6%) |
| Clonic | 39 (10.1%) |
| Epileptic spasms | 31 (8.0%) |
| Myoclonic | 44 (11.4%) |
| Tonic | 66 (17.1%) |
| Infantile spasms | 116 (30.1%) |
| --- SEIZURE TYPES: FOCAL --- |  |
| Absence | 45 (11.7%) |
| Focal | 97 (25.1%) |
| --- DEVELOPMENTAL QUOTIENT --- |  |
| Developmental Quotient (overall) | 25.7 (9.6–42.3) |
| Developmental Quotient — Fine Motor | 5.8 (5.0–35.0) |
| Developmental Quotient — Gross Motor | 13.3 (8.3–25.0) |
| Developmental Quotient — Social/Communication | 28.3 (5.6–50.0) |
| Developmental Quotient — Language | 16.7 (5.0–89.4) |
| --- DEVELOPMENTAL DELAY SEVERITY CATEGORIES --- | 342 |
| Profound Developmental Delay (0–30) | 190 (55.6%) |
| Severe Developmental Delay (30–50) | 88 (25.7%) |
| Moderate Developmental Delay (50–70) | 50 (14.6%) |
| Mild Developmental Delay/Neurotypical (70+) | 14 (4.1%) |

**Supplementary Table 2**. Enrichment analysis results for pathogenic versus benign variants across SCN8A segments. *See Excel file.*

**Supplementary Table 3.** Enrichment of features within 50 or 100 nucleotides of post-translational modification sites, binding sites, disordered regions, and structural sites. *See Excel file.*

**Supplementary Table 4.** Bootstrapped confidence intervals for enrichment of features in variants predicted to be splice-altering by individual tools. *See Excel file.*

**Supplementary Table 5.** Correlation matrix of features with recurrent variants, variants within 50 nucleotides of post-translational modification sites and structural sites, and variants that are likely splice-altering by the composite metric. Values are reported as R^2^ (p). *: p<0.05, ** p<0.01. *See Excel file.*
